# Household transmission of SARS-CoV-2 from humans to pets in Washington and Idaho: burden and risk factors

**DOI:** 10.1101/2021.04.24.440952

**Authors:** Julianne Meisner, Timothy V. Baszler, Kathryn H. Kuehl, Vickie Ramirez, Anna Baines, Lauren A. Frisbie, Eric T. Lofgren, David M. DeAvila, Rebecca M. Wolking, Dan S. Bradway, Hannah Wilson, Beth Lipton, Vance Kawakami, Peter M. Rabinowitz

## Abstract

SARS-CoV-2 is believed to have emerged from an animal reservoir; however, the frequency of and risk factors for inter-species transmission remain unclear. We carried out a community-based study of pets in households with one or more confirmed SARS-CoV-2 infection in humans. Among 119 dogs and 57 cats with completed surveys, clinical signs consistent with SARS-CoV-2 were reported in 20 dogs (21%) and 19 cats (39%). Out of 81 dogs and 32 cats sampled for testing, 40% of dogs and 43% of cats were seropositive, and 5% of dogs and 8% of cats were PCR positive; this discordance may be due to delays in sampling. Respondents commonly reported close human-animal contact and willingness to take measures to prevent transmission to their pets. Reported preventative measures showed a slightly protective trend for both illness and seropositivity in pets, while sharing of beds and bowls had slight harmful effects.

## Background

Coronaviruses infect multiple mammalian species, and SARS-CoV-2 virus, the etiological agent of COVID-19 infection, likely jumped to humans from a mammalian source [1]. While currently the virus is spreading person to person, the ACE2 receptor involved in SARS-CoV-2 transmission is present in multiple species and there are numerous reports of infections in pets [2–4]. Currently, 110 domestic cats and 95 domestic dogs in the USA have been reported by USDA-APHIS to have SARS-CoV-2 infection [5]. Workplace transmission of SARS-CoV-2 between humans and animals has also been documented, including in zoos (felids and non-human primates) and on mink farms [6,7]. This is consistent with previous reports of SARS-CoV-1 infecting cats and ferrets, and laboratory studies demonstrating experimental SARS-CoV-2 infection of non-human primates, ferrets, hamsters, and rabbits [8]. Less is known, however, about the frequency of and risk factors for SARS-CoV-2 transmission between humans and companion animals in a household setting. Furthermore, the natural history of SARS-CoV-2 infection in pets is poorly understood.

Given the close contact many people have with their pets and the intimate nature of their shared environment, exacerbated during periods of human quarantine or isolation, it is important to better understand the role of companion animals in community infection patterns, including contribution to virus evolution and emergence of novel strains. In light of evidence from mink farms that animal-origin variants may contain spike mutations and other changes that could affect clinical features of infection [9,10], recent evidence suggesting mouse origins of the omicron variant [11], and Hong Kong’s recent decision to cull 2,000 hamsters after a pet shop worker was infected with the delta variant [12], ongoing monitoring of SARS-CoV-2 transmission between humans and animals in household and other human-animal contact settings remains critical.

We report our findings from the COVID-19 and Pets Study (CAPS), an ongoing cross-sectional community-based study of pets in households of persons with documented COVID-19 infection. The goal of the study is to describe the frequency of transmission between humans and animals within a household, and to determine human, animal, and environmental risk factors for that transmission, in a One Health framework.

## Methods

The COHERE [13] and STROBE [14] statements were used to guide reporting of the findings and the preparation of this manuscript.

### Study population

We defined a household as one or more persons ages 18 or older, co-housing with at least one pet that does not live solely outdoors. Pets were defined as dogs, cats, ferrets, and hamsters based on prior research documenting experimental COVID-19 infection in these species [15,16].

We conducted this study in King, Snohomish, Yakima, Whitman, Pierce, Spokane, and Benton counties in Washington, and Latah County in Idaho. This paper reports on sampling conducted from April 2020 to September 2021.

### Study design

CAPS is a cross-sectional study with individual- and household-level data collection. Study participation involved two components, detailed below: an online survey followed by animal sampling.

### Recruitment and eligibility

Households were recruited through partnerships with other COVID-19 clinical trials and community studies, social media, word of mouth, community partners, and by contact tracers from Public Health—Seattle & King County during case investigation/contact tracing calls. Individuals were screened for eligibility using the UW Research Electronic Data Capture (REDCap) system [17], a HIPAA-compliant web tool for clinical research, with criteria including county of residence, pet ownership, and one or more household member with confirmed SARS-CoV-2 infection via PCR or antigen testing by a provider or laboratory. Animals with known fearful and/or aggressive behavior were excluded, however, other animals in the corresponding household were eligible.

### Ethical approvals

This study and its protocols received ethical approval from the University of Washington’s Institutional Review Board STUDY00010585) and Office of Animal Welfare (PROTO201600308: 4355-01). Informed consent was obtained via REDCap, or over the phone with the study coordinator, after the nature and possible consequences of study involvement had been explained. Once eligibility was confirmed and consent was obtained, individuals completed the online survey.

### Survey

A comprehensive survey was completed by a household member prior to scheduling of the sampling visit. Surveys were completed online by the study participant using the REDCap interface, or via phone with the study coordinator. Human items included COVID-19 symptom onset, specific symptoms experienced and severity; comorbidities; vaccination status including dates and type; and reported COVID-19-like illness of any other household members, including those who did not have confirmatory testing. Animal items, stratified on individual animal, included veterinary clinical variables, history of illness compatible with SARS-CoV-2 infection, and contact between individual animals and individual members of the household. Environmental items included type and size of home, type of flooring (carpet, wood, etc.), and availability of outdoor space for pets to roam.

A second survey was completed verbally at the time of sampling on any changes in the clinical status of human and animal household members since the REDCap survey was completed, including new hospitalizations, symptoms, or COVID-19 diagnoses. Confirmation of SARS-CoV-2 test date and positive result was also performed through review of test results by the sampling team. Self-test results were not accepted.

### Animal sampling

Sampling was performed by a team of two study personnel including at least one veterinarian. In most cases sampling was conducted at the participant’s home; however, several animals were tested at veterinary hospitals. No chemical restraint was used, nor muzzles due to biosafety concerns.

Species-appropriate restraint was employed using standard techniques to allow for venipuncture and collection of 3 mL of blood into a labeled serum separator tube. Following venipuncture, swab samples were collected from both rostral nares/nasal passage and the caudal oropharynx and then placed into one Primestore Molecular Transport Medium (MTM) tube. A separate fecal swab was collected from the rectum and placed into a separate Primestore MTM tube. All participants received educational information from the field team about measures to mitigate household COVID-19 transmission.

Swab and serum samples were transported on ice within 24 hours to the Washington Animal Disease Diagnostic Laboratory (WADDL) for PCR and antibody testing.

### Testing

#### SARS-CoV-2 RT-PCR

Respiratory or fecal swabs: RNA extraction and SARS-Cov-2 reverse transcriptase (RT) real-time PCR was performed as described [18]. Following initial viral detection by PCR, three dog samples and one cat sample were submitted to University of Minnesota Genomics Center (Oakdale, MN 55128) for whole genome sequencing (WGS) [19]. A second cat sample was submitted to the USDA National Veterinary Services Laboratories (NVSL) in Ames Iowa for WGS. Mutational analysis was performed using the GISAID EpiFlu Database CoVsurver: Mutation Analysis of hCoV-19 [20,21]. All five sequences were deposited into GISAID, with accession numbers EPI_ISL_7845315, EPI_ISL_7845316, EPI_ISL_7845317, EPI_ISL_7845318, and EPI_ISL_8897004. SARS-CoV-2 lineages were assigned using the Phylogenetic Assignment of Named Global Outbreak LINeages (Pango lineage) tool [22,23].

#### SARS-CoV-2 Spike Protein Receptor Binding Domain (RBD) ELISA

WADDL developed canine and feline SARS-CoV-2 ELISA assays using recombinant SARS-CoV-2 Spike Receptor Binding Domain protein as antigen (S-RBD). The recombinant RBD was obtained from the UW Center for Emerging and Reemerging Infectious Disease (CERID) laboratory of Dr. Wesley Van Voorhis through an institutional Material Transfer Agreement. WADDL used an in-house standard operating procedure for indirect ELISA of SARS-CoV-2 in 96-well format based on a previous publication in humans [24]. The major components of the assay included: 1) rS-RBD coating of plates as target antigen (2ug/ml in Sigma Carbonate-Bicarbonate Buffer); 2) 1:100 dilution of test sera (diluted in ChronBlock ELISA Buffer-Chondrex Inc.); 3) anti dog IgG-HRP as linker (Southern BioTech goat anti-canine IgG) and 4) Sigma (TMB) liquid substrate system to develop OD. Plates were blocked with ChronBlock ELISA buffer per manufacturer’s instructions, washing solution consisted of PBS+0.1% Tween 20 (Sigma), and plates were read on a plate reader at 450 nM. Test sera were run in triplicate and utilized at “*test OD*”.

For the canine RBD ELISA, the negative controls consisted of sera from six pre-COVID dogs, archived at WADDL and tested for canine Adenovirus (CAV), canine Distemper Virus (CDV), canine Coronavirus (CCV), canine Parainfluenza (CPI), and canine Parvovirus (CPV) IgG. All six samples had antibody presence of one or more of the tests performed, however no sera reacted in the SARS-CoV-2 canine RBD ELISA. For the cat RBD ELISA, the negative controls consisted of sera from three pre-COVID cats from WADDL archives, tested for feline Coronavirus (FIP-FeCV) and feline Panleukopenia Virus (FPV)-IgG. Two of the three samples had antibody presence of one or more of the tests performed (including 2 for FIP-FeCV); however, neither reacted in the SARS-CoV-2 feline RBD ELISA. Negative controls were run in triplicate and the mean was utilized as “*negative control OD*” A ratio of *test OD: negative control OD* was used to determine the results. The positive cutoff of 2.0 *test OD: negative control OD* ratio equated to the mean of negative controls + 3 standard deviations of the mean.

SARS-CoV-2 RBD ELISA was repeated three times for all samples, and the final results were tabulated as a mean value obtained from the repeated testing. As no dog or cat in Washington or Idaho had been confirmed to be SARS CoV-2 positive via serology prior to our study, the first antibody positive case for each species and state was sent to the NVSL for confirmation via virus neutralization (VN) assay in keeping with regulatory recommendations. Both canine and feline SARS-CoV-2 RBD ELISA positive samples were confirmed at NVSL by VN.

### Statistical analyses

The primary aim of this study was to estimate the burden of household SARS-CoV-2 transmission from humans to their pets. Secondary aims included describing the nature of human-animal contact within households and identifying risk factors for household transmission, including human-animal contact. All analyses were conducted in R [25].

#### Outcome

Animal infection with SARS-CoV-2 was defined as an animal meeting one or more of the following criteria: (1) SARS CoV-2 RBD ELISA seropositive status, (2) PCR positive status, or (3) illness consistent with SARS-CoV-2 infection, hereafter referred to as “illness,” defined as participant answer of “yes” to the survey question: “Since the time of COVID diagnosis/symptom onset in the household, has this animal had any new issues with difficulty breathing, coughing and/or decreased interest in playing, walking, or eating?” Serostatus was parameterized as ELISA ratio, log-transformed for the sake of interpretability; PCR positive status and illness were parameterized as binary variables.

#### Regression models

Outcome was defined as an animal case of SARS-CoV-2 (definition above). Separate regression models were fit for each outcome definition.

Household-level exposures for animal infection included residence in house versus apartment or condominium (binary), home size in square feet (continuous), and the number of human confirmed SARS-CoV-2 cases (continuous). Animal-level exposures for infection included bedsharing with one or more human household members (binary), sharing bowls with one or more household members (binary), and SARS-CoV-2 positive household members taking precautions to prevent transmission to their pets following diagnosis, including not petting or kissing the animal, staying in a different room, and having someone else feed and walk the animal (binary). We also examined the association between canine seropositivity and illness compatible with SARS-CoV-2 infection in the animal, and between seropositivity and time since the animal was first exposed, defined as two days prior to the first date any household member had symptoms of COVID-19 or tested positive, whichever was earlier.

We identified possible confounders *a priori* using a directed acyclic graph (DAG; Figure 1). The minimum sufficient adjustment set was defined, using this DAG and DAGitty.net, separately for each exposure [26]. Animal species was explored as an effect modifier using a multiplicative interaction term, and stratified results presented in all cases in which this interaction term reached statistical significance (p ≤ 0.05).

**Figure 1:**
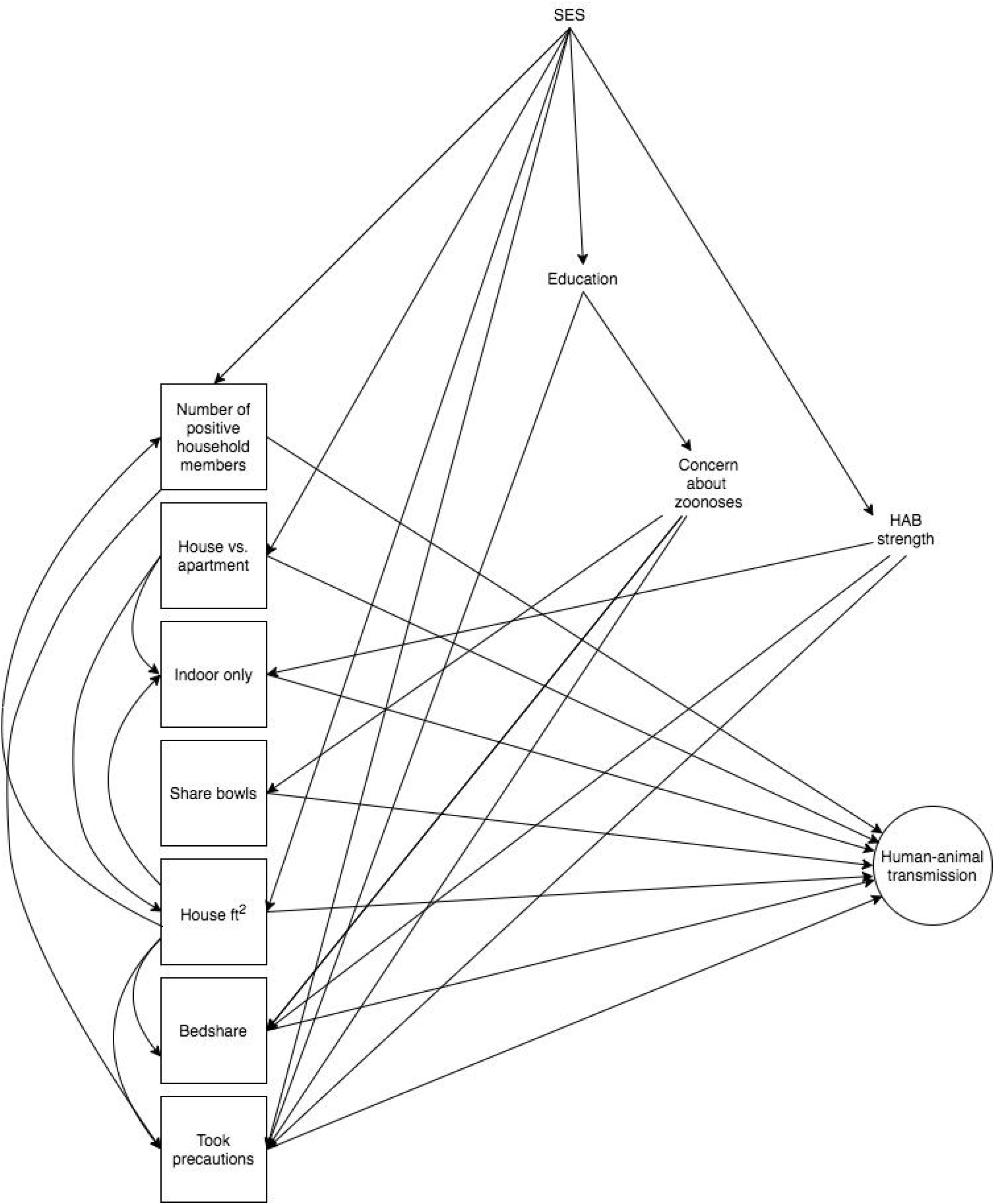
Directed acyclic graph for human-animal SARS CoV2 transmission. Variables outlined with a square are the exposures of interest, while outcome (approximated by serostatus, PCR result, and illness in separate models) is outlined with a circle. HAB: human-animal bond; SES: socioeconomic status; took precautions: SARS-CoV-2 positive household member(s) took precautions to prevent transmission to pet; indoor-only: animal does not go outdoors; bedshare: animal shares a bed with one or more household members.

For each exposure of interest we implemented a generalized estimating equation (GEE) approach with an exchangeable working correlation structure, household as the clustering variable, and binomial models with a logit (binary outcomes) or Gaussian (continuous outcomes) link, using the geepack package in R [27]. For regression of ELISA ratio on illness and time since first exposure, we performed linear regression using the glm() function in R.

## Results

### Recruitment

In total, 107 eligible households enrolled and completed the survey. No households currently living as unhoused enrolled. Two households corresponded to a single dog which was moved from the participant’s home to a family member’s home immediately after the onset of the participant’s COVID-19 symptoms, leaving 105 households corresponding to 119 dogs and 57 cats available for analyses; no ferrets or hamsters enrolled or were sampled.

Sample collection is detailed in Figure 2. In total, 83 households corresponding to 100 dogs and 47 cats had a sampling visit conducted. Of these, six dogs and eight cats belonged to households were not sampled due to temperament, leaving 94 dogs and 39 cats with PCR results, while an additional 13 dogs and 9 cats were safe to restrain for swab (PCR) samples but not for serum collection, leaving 81 dogs and 32 cats with serology results.

**Figure 2:**
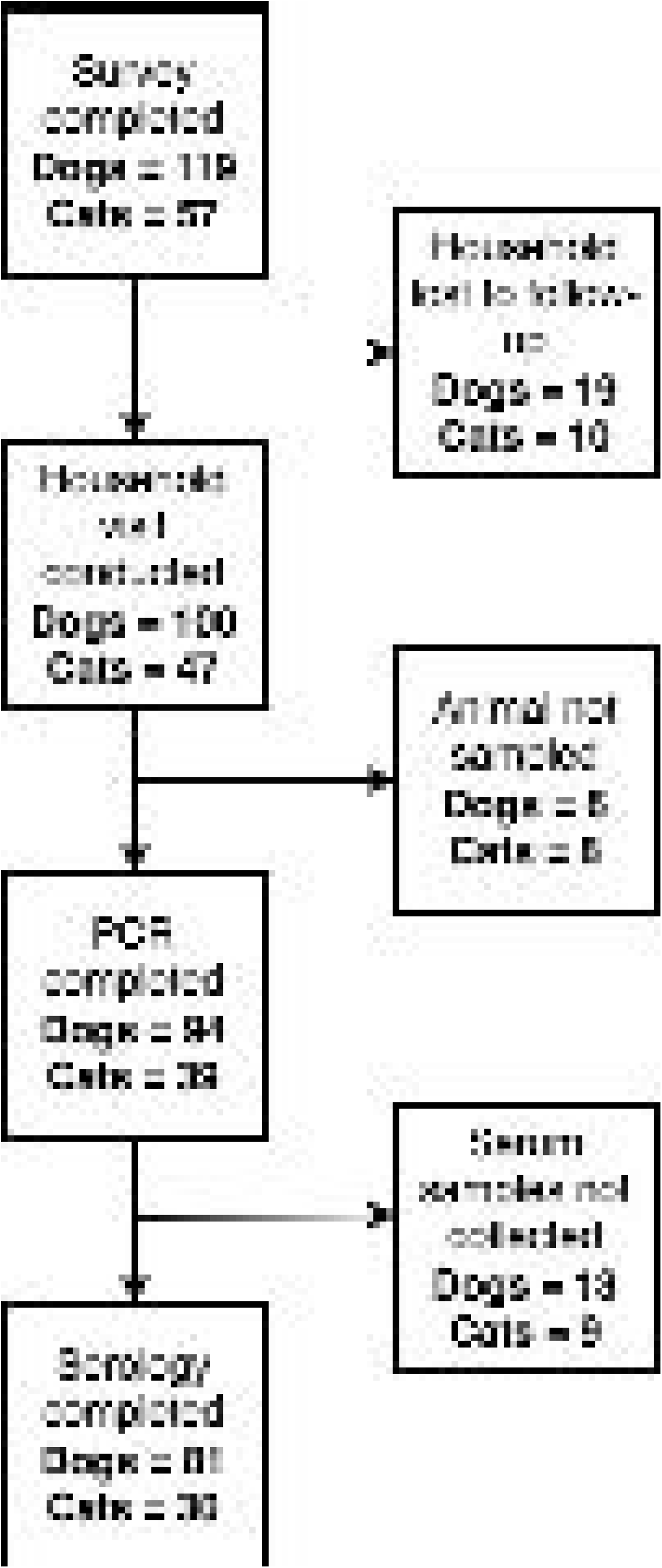
Flowchart depicting serological and PCR sampling. Out of 119 dogs and 57 cats corresponding to 105 households with completed surveys, PCR testing is complete for 94 dogs and 39 cats, and serological testing is complete for 81 dogs and 32 cats. The remaining pets were not sampled due to safety concerns.

### Descriptive statistics

Descriptive statistics are presented in Table 1. On average, at least six weeks (dogs) and two weeks (cats) elapsed between the last human COVID-19 diagnosis in the household and animal sampling. Of the 119 dogs and 57 cats with completed surveys, 20.4% (95% CI 12.9%, 29.7%) of dogs and 38.8% (95% CI 25.2%, 53.8%) of cats had reported illness. Of the 94 dogs and 39 cats who were PCR tested, 5.3% (95% CI 1.8%, 12%) of dogs and 7.7% (95% CI 1.6%, 20.9%) of cats were positive; of the 81 dogs and 32 cats who had serum collected, 40.2% (95% CI 29.6%, 51.7%) of dogs and 40.6% (95% CI 23.7%, 59.4%) of cats were seropositive. Individual animal SARS-CoV-2 RBD ELISA results are shown in Figure 3 (dogs) and Figure 4 (cats). SARS-CoV-2 RBD ELISA test OD:negative control OD ratios in seropositive animals ranged from 2.03 – 21.22 in dogs and from 3.01 – 30.35 in cats.

**Figure 3:**
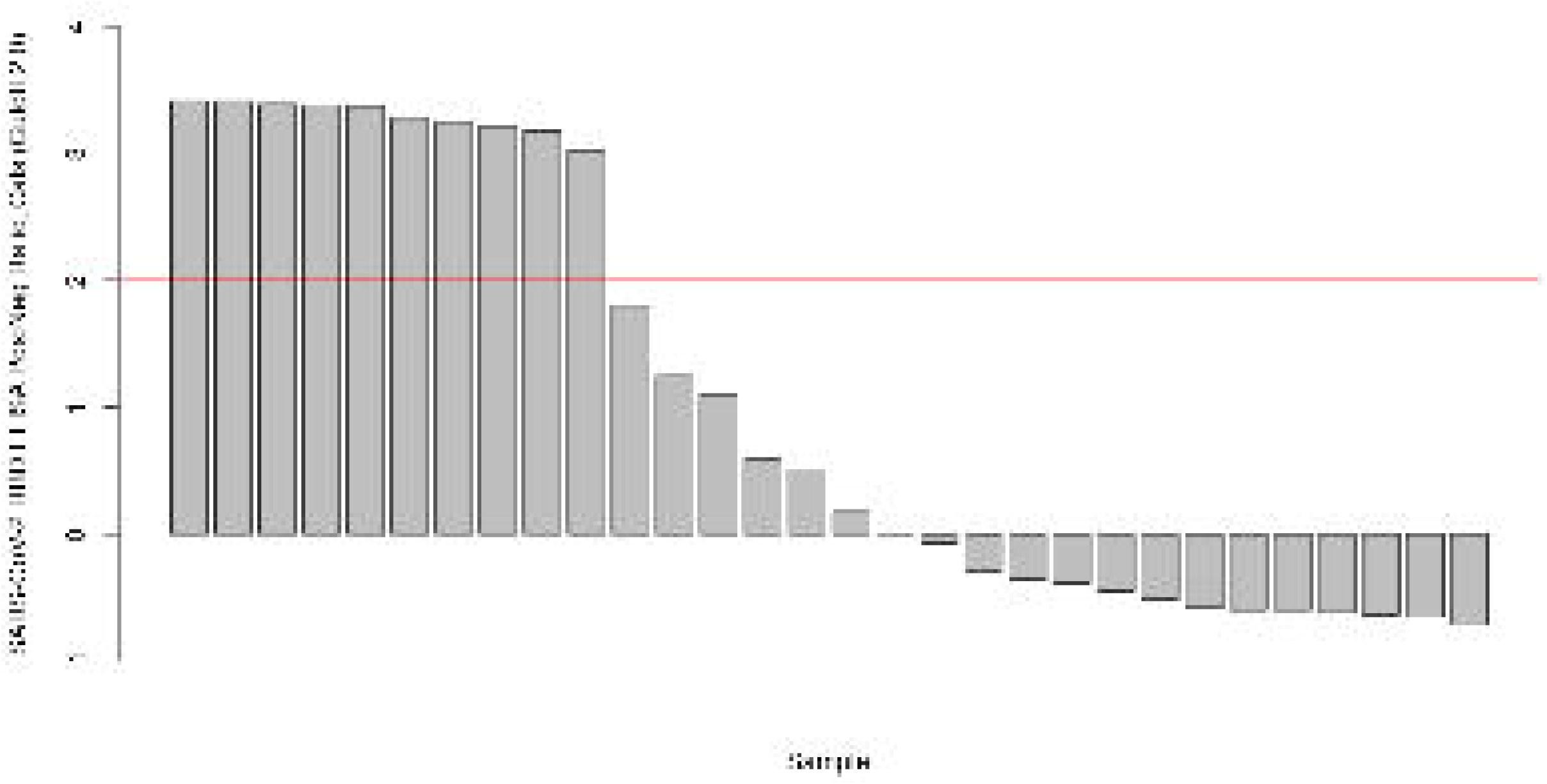
SARS-CoV-2 RBD ELISA Serology data, cats. PCR testing is complete for 39 cats, and serological testing is complete for 32 cats. The remaining pets were not sampled due to safety concerns.

**Figure 4:**
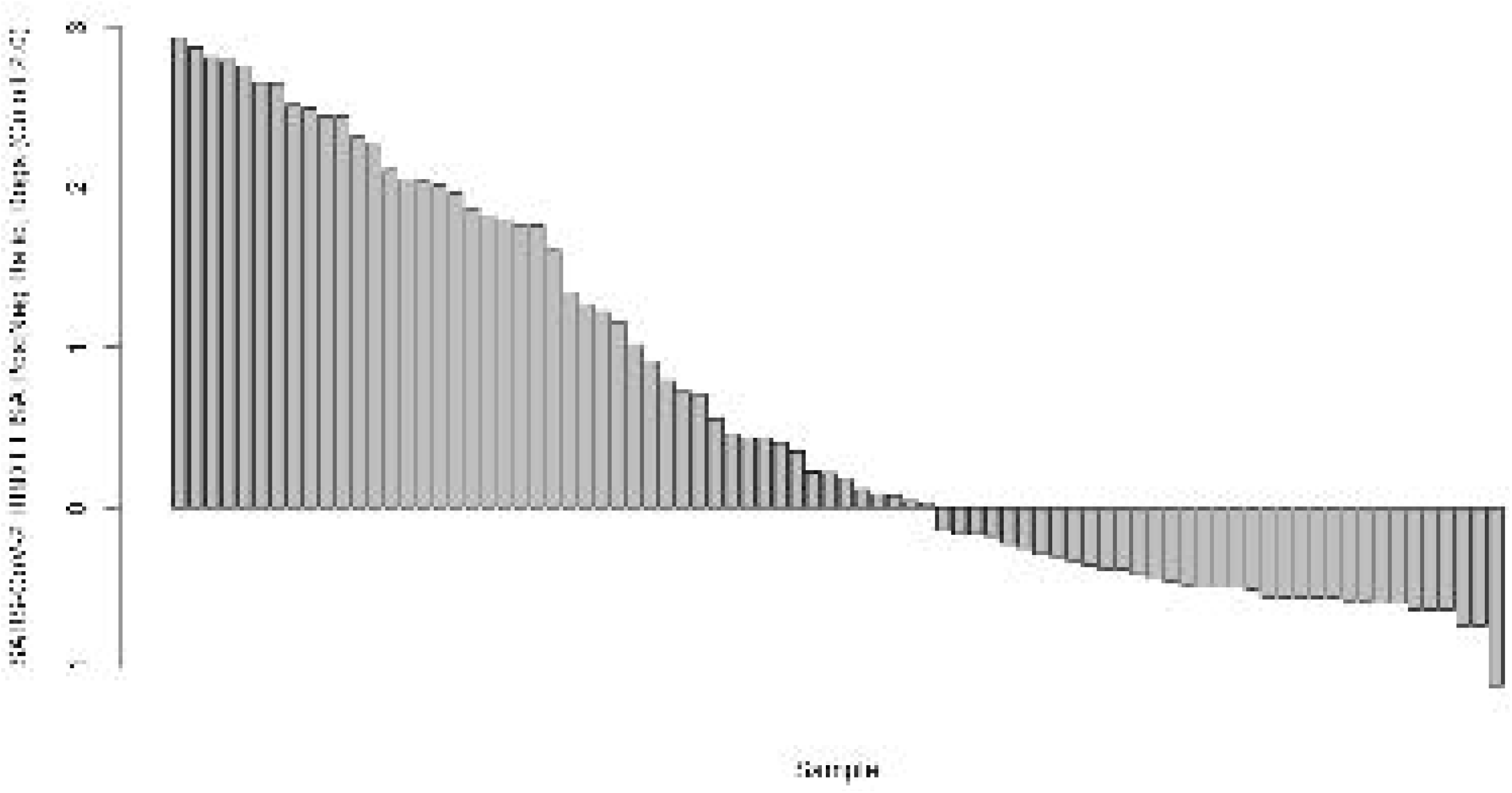
SARS-CoV-2 RBD ELISA Serology data, dogs. PCR testing is complete for 94 dogs, and serological testing is complete for 81 dogs. The remaining pets were not sampled due to safety concerns.

**Table 1:**
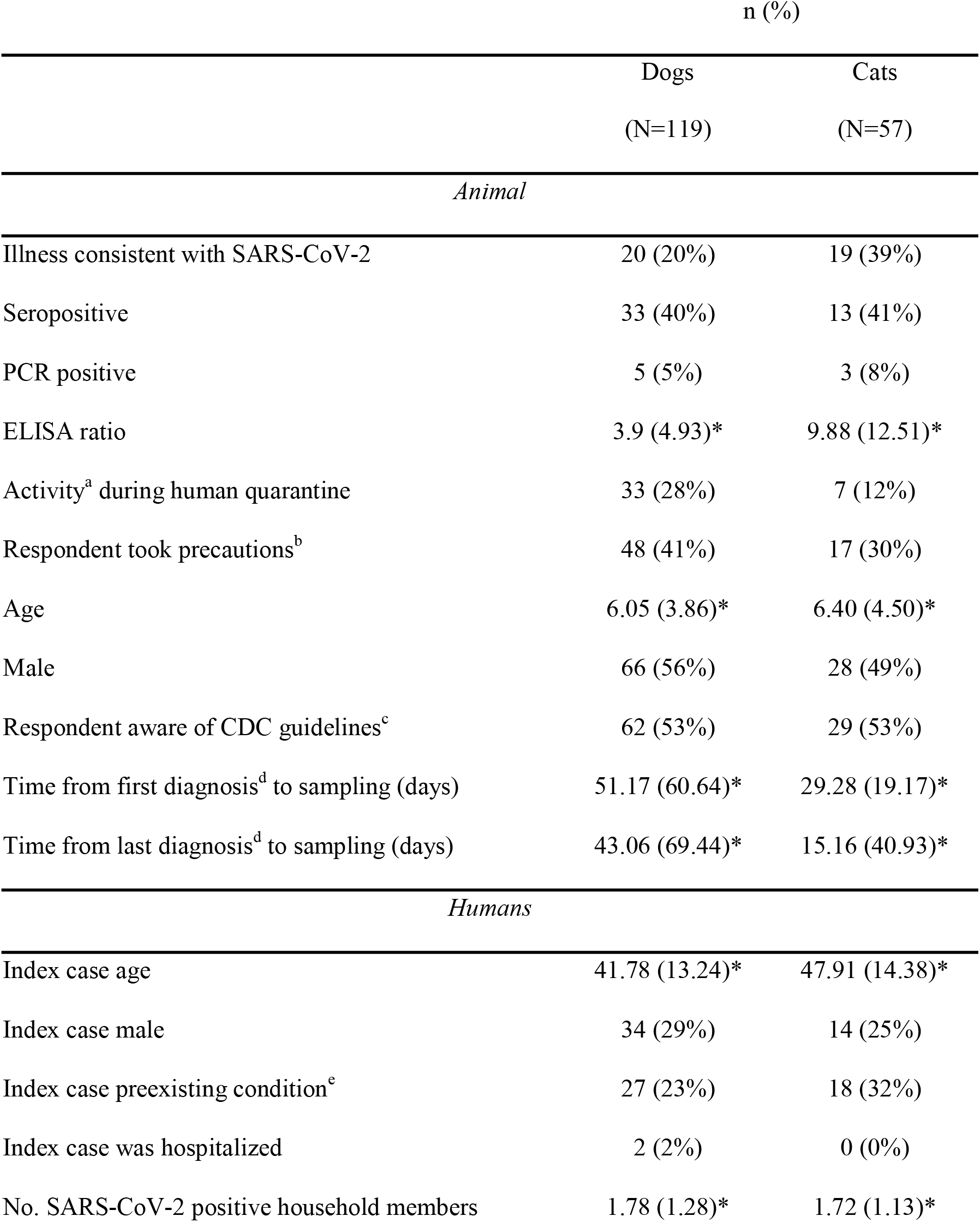
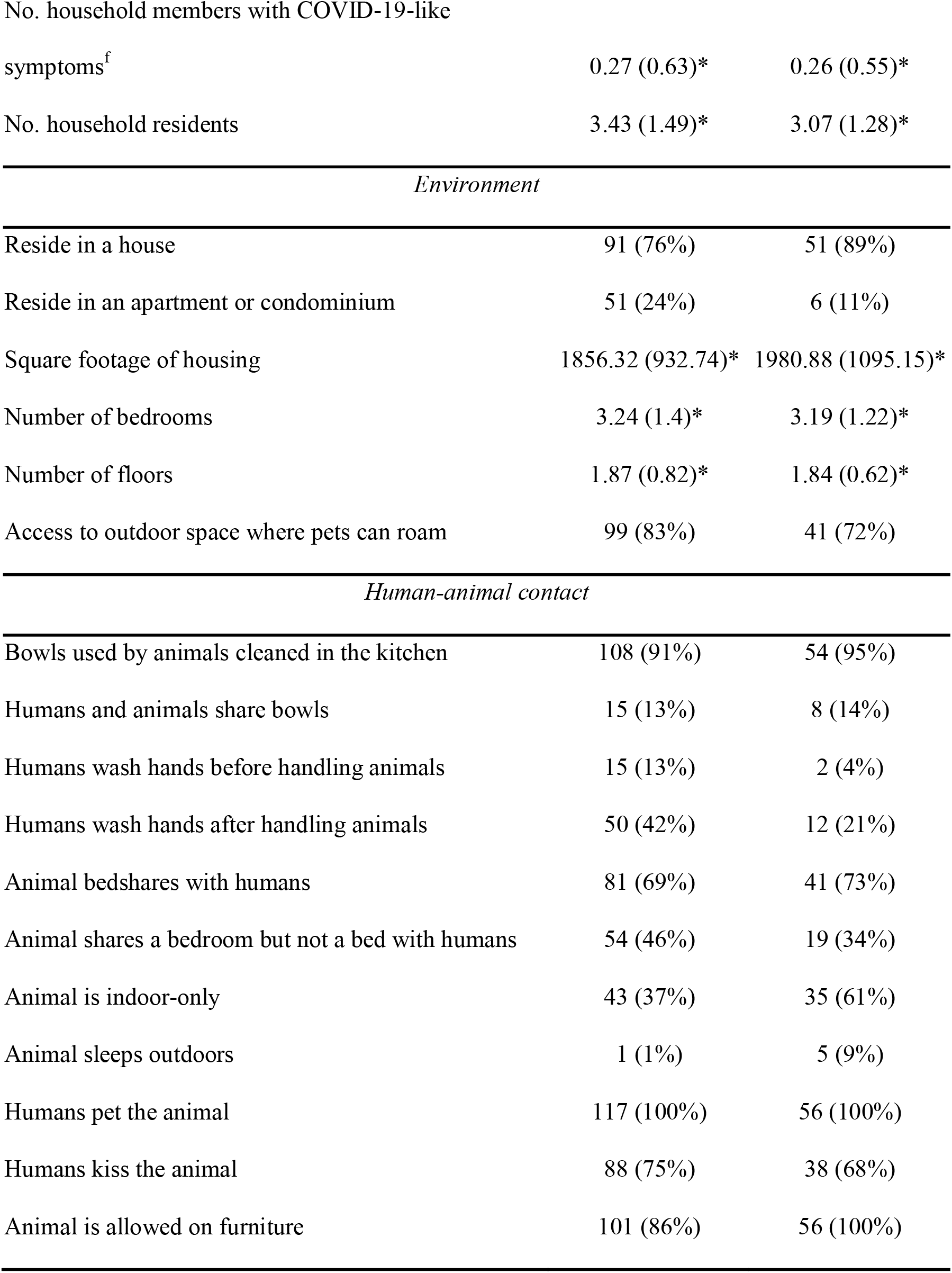
Descriptive statistics for 119 dogs and 57 cats corresponding to 105 households. *mean (standard deviation). ^a^Activity defined as going to a veterinary clinic or groomer, being walked off-leash, or visiting an off-leash park, dog park, kennel, or daycare facility. ^b^Precautions to prevent human-animal SARS-CoV-2 transmission following diagnosis: not petting or kissing the animal, staying in a different room, and having someone else feed and walk the animal. ^c^Guidelines to prevent human-animal SARS-CoV-2 transmission. ^d^First diagnosis: earliest known, confirmed SARS-CoV-2 diagnosis in the household; final diagnosis: last known, confirmed SARS-CoV-2 diagnosis in the household. ^e^Prexisting conditions: diabetes, kidney disease, heart disease, hypertension, immunosuppression. ^f^Household members who had COVID-19-like symptoms but did not get tested.

Five dog swabs (Cts 26.08 – 37.67) and 3 cats (Cts 27.03 – 39.97) were PCR positive on nasal/oropharyngeal swabs; one of these dogs was also PCR positive from fecal swab (Ct 39.20). Five PCR positive samples (2 cats and 3 dogs) had Cts sufficient for WGS (Ct<30): The earliest cat sample (April 2021) that underwent WGS fell into Pango clade B.1.2. A later dog sample sequenced as Delta sublineage B.1.617.2.103 (AY103), while the other three (2 cat, 1 dog) samples sequenced as Delta sublineage B.1.617.2.25 (AY25). Of the five PCR positive dogs, three were PCR positive prior to being seropositive and two were simultaneously PCR and seropositive.

There were 11 households with two or more positive animals, and among multi-pet households with at least one positive pet, mean prevalence (PCR or serology) was 91%. Out of eight total PCR positive cases, all were detected after April 2021, when the first case of the Delta variant was documented in Washington State.

Nearly one-third of dogs engaged in activities outside of the household during periods of human isolation or quarantine. Over 50% of both cats and dogs resided in households whose residents reported awareness of CDC guidelines to prevent human-animal transmission of SARS-CoV-2, and 48 (41%) dogs and 17 (30%) cats resided in households which reported taking precautions to prevent such transmission to household pet(s) following diagnosis. No cats and only two dogs resided in a household in which an infected person was hospitalized for COVID-19. Nearly all dogs (83%) and most cats (72%) had access to yards or gardens and were allowed on furniture (86% of dogs and 100% of cats), and the majority were kissed by (75% of dogs and 68% of cats) and shared beds (69% of dogs and 73% of cats) with human household members. Almost all dogs’ (91%) and cats’ (95%) bowls were washed in the kitchen.

### Regression models

Results of regression models are presented in Table 2 as prevalence odds ratios for the binary outcome of illness, reflecting the cross-sectional design of this study, and as exp^β^ for the outcome of ELISA ratio, which can be interpreted as the relative change (ratio scale) in ELISA ratio for a one unit change in the exposure. As so few animals were PCR positive, we did not run regression models for this outcome. With the exception of house size, which was adjusted for house type as the minimum sufficient adjustment set was very small for this exposure, confounders were not adjusted for due to concerns regarding overfitting arising from the small sample size. Effect modification by species was found only for house type.

**Table 2:**
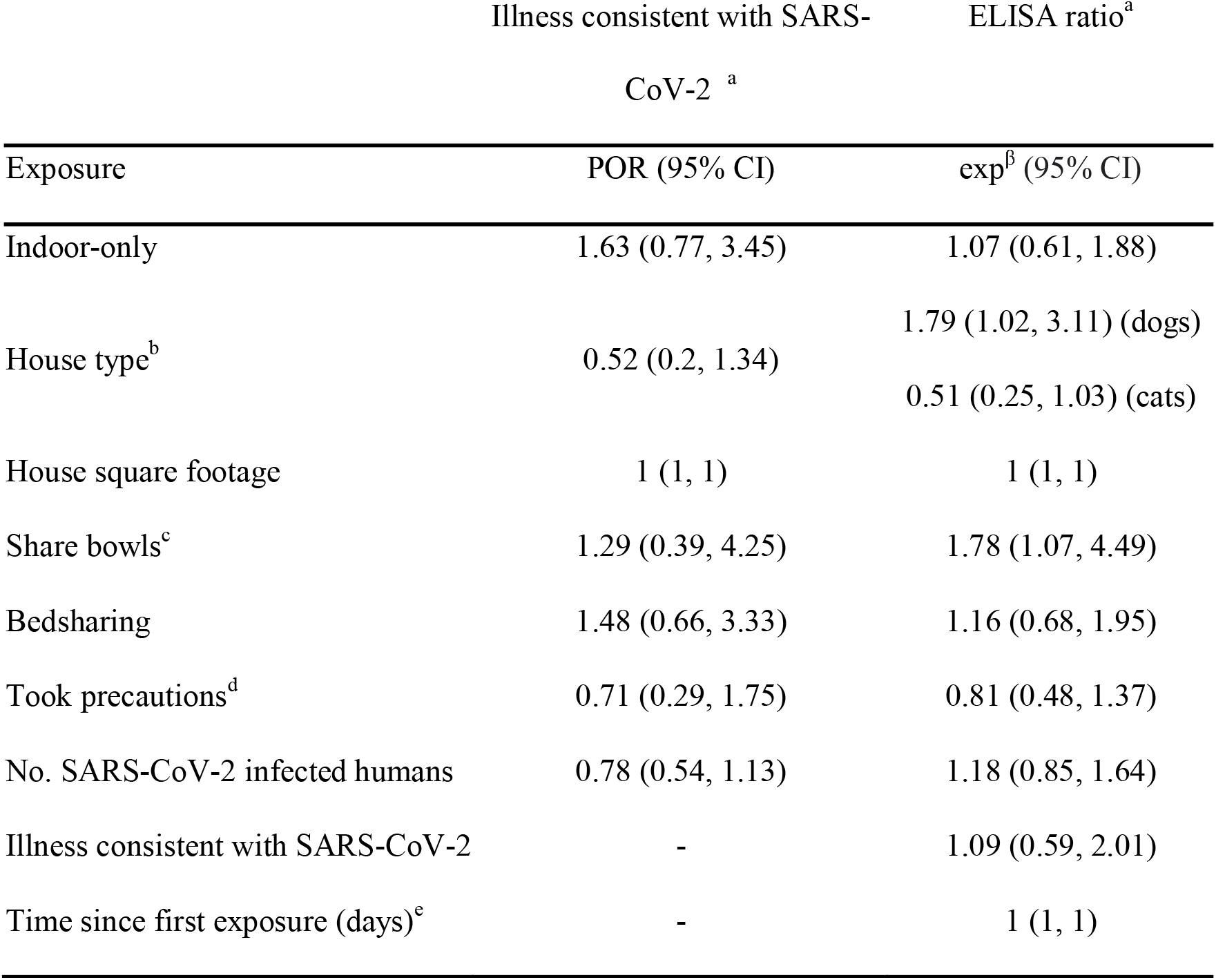
Regression model results. House size was adjusted for house type, but no other models were not adjusted for confounders due to overfitting concerns. ^a^Survey results available for 119 dogs and 57 cats, serology results available for 81 dogs and 32 cats. ^b^House versus apartment or condominium. ^c^Animals and humans share bowls. ^d^Precautions taken to prevent human-animal SARS-CoV-2 transmission following diagnosis: not petting or kissing the animal, staying in a different room, and having someone else feed and walk the animal. ^e^First exposure defined as 2 days prior to first positive diagnosis in the household or onset of symptoms, whichever was earlier. POR: prevalence odds ratio; 95% CI: 95% confidence interval.

Dogs residing in houses on average had a 79% (95% CI 2%, 211%) higher ELISA ratio than dogs residing in apartments or condos, while the inverse association was detected for cats (49% lower mean ELISA ratio, 95% CI 75% lower, 3% higher) and for the outcome of illness in both cats and dogs (48% lower prevalence odds, 95% CI 80% lower, 34% higher); this association reached statistical significance for dogs only. No other effect estimates reached statistical significance; however, there were positive trends across both outcome definitions for bed sharing with humans, sharing bowls, and being indoor only; and a negative effect for precautions taken to prevent SARS-CoV-2 transmission following diagnosis. We also found ELISA ratio was positively associated with illness; however, we did not find evidence of an effect of time since first exposure on ELISA ratio, nor of house square footage on either outcome.

## Discussion

We present the results of a cross-sectional, One Health study of SARS-CoV-2 transmission between people and their pets. The study results indicate that household transmission of SARS-CoV-2 from humans to animals occurs frequently and infected animals commonly display signs of illness. Notably, in 9 out of 11 households with multiple pets of whom at least one tested positive (PCR or serology), all tested pets were positive. We furthermore show that close human-animal contact is common among people and their pets in this study population, that this contact appears to facilitate SARS-CoV-2 transmission, and that pet owners are familiar with and willing to adopt measures to protect their pets from COVID-19.

There are several limitations to our approach. First, several weeks had elapsed from first reported exposure to household sample collection from animals in most households, possibly limiting our ability to detect viral shedding by PCR testing but strengthening our ability to detect seroconversion. Second, while we assume transmission is from humans to pets, the cross-sectional nature of this study precludes certainty regarding the direction of transmission. Nevertheless, as SARS-CoV-2 is transmitted predominantly human-to-human, few cases of SARS-CoV-2 have been documented in dogs and cats, and no cases have been documented to be transmitted from dogs or cats to humans, we believe transmission in this study was exclusively from humans to pets. Third, our study is subject to residual confounding due to inability to adjust for confounders without risking over-fitting. We do not expect unmeasured or unadjusted confounders to exert strong effects other than latent (and therefore difficult to measure and model) constructs, such as socioeconomic status, strength of the human-animal bond, and level of concern about zoonotic disease transmission. Finally, our definition of illness in pets is simple, derived from a single survey item, and vulnerable to misclassification if these clinical signs are due to other etiologies. This survey was created early in the COVID-19 pandemic, although illness in pets is still not well-characterized.

We believe respondents misunderstood the question, “Is this animal indoor only vs. indoor/outdoor” as 37% of dogs were reported to be indoor-only, however we believe this variable retains its connection to degree of animal contact despite mismeasurement (i.e., a dog labeled as “indoor only” likely spends more time in an indoor setting with humans than a dog labeled as indoor/outdoor). We do not expect strong measurement error in any of the other variables examined. As no gold-standard for canine anti-SARS-CoV-2 serology exists, validation of our ELISA assay was limited to analytic validation and we could not reliably estimate diagnostic sensitivity of our serological test; full diagnostic validation was not possible due to the absence of sufficient gold-standard positive and negative samples, a limitation arising from the status of SARS-CoV-2 as an emerging pathogen. However, all pre-COVID-19 samples evaluated were negative, indicating specificity approaches 100%, and all samples sent to USDA-NVSL for confirmatory PCR and serology testing had concordant results. While our primary aim—to estimate the burden of human-animal SARS-CoV-2 transmission—was estimated with reasonable precision, due to our small sample size variance was high for effect estimates produced by our regression model. Finally, by nature of our recruitment methods and study population, generalizability of our findings is likely limited to highly-educated, higher-income individuals residing in urban and suburban communities.

## Conclusions

These limitations aside, our study contributes important and novel findings to the literature on cross-species transmission of SARS-CoV-2, with relevance to other zoonoses transmitted in a household setting. Furthermore, we collected human, animal, and environmental data, representing a true One Health approach to this critical research question. Finally, our findings indicate households in this population are willing to adopt measures to protect their pets from SARS-CoV-2 infection, and that these measures may be effective, indicating an opportunity to prevent household transmission of zoonoses through health education and policy.

## Acknowledgements

Data collection: Jessica Bell, DVM and Raelynn Farnsworth, DVM, Washington State University College of Veterinary Medicine; Katherine Burr, DVM and Gemina Garland Lewis, MS, Center for One Health Research, University of Washington.

Survey Review: J. Scott Weese, Ontario Veterinary College, University of Guelph.

Recombinant SARS-CoV-2 receptor binding domain source material: Dr. Wes Van Voorhis, Center for Emerging and Re-emerging Infectious Diseases, University of Washington, Seattle, WA, USA [28,29].

## Author Bio

Julianne Meisner is a post-doctoral fellow in Environmental and Occupational Health Sciences at the University of Washington. Her research interests lie at the intersection of human and animal health, and the application of rigorous epidemiologic methods to research at this interface.

